# GWAS on family history of Alzheimer’s disease

**DOI:** 10.1101/246223

**Authors:** Riccardo E. Marioni, Sarah E. Harris, Allan F. McRae, Qian Zhang, Saskia P. Hagenaars, W. David Hill, Gail Davies, Craig W. Ritchie, Catharine Gale, John M. Starr, Alison M. Goate, David J. Porteous, Jian Yang, Kathryn L. Evans, Ian J. Deary, Naomi R. Wray, Peter M. Visscher

## Abstract

Alzheimer’s disease (AD) is a public health priority for the 21^st^ century. Risk reduction currently revolves around lifestyle changes with much research trying to elucidate the biological underpinnings. Using self-report of parental history of Alzheimer’s dementia for case ascertainment in a genome-wide association study of over 300,000 participants from UK Biobank (32,222 maternal cases, 16,613 paternal cases) and meta-analysing with published consortium data (n=74,046 with 25,580 cases across the discovery and replication analyses), six new AD-associated loci (P<5x10^−8^) are identified. Three contain genes relevant for AD and neurodegeneration: *ADAM10, ADAMTS4,* and *ACE*. Suggestive loci include drug targets such as *VKORC1* (warfarin dose) and *BZRAP1* (benzodiazepine receptor). We report evidence that association of SNPs and AD at the *PVR* gene is potentially mediated by both gene expression and DNA methylation in the prefrontal cortex. Our discovered loci may help to elucidate the biological mechanisms underlying AD and, given that many are existing drug targets for other diseases and disorders, warrant further exploration for potential precision medicine applications.

## Introduction

The social and economic impact of Alzheimer’s disease (AD) makes it a global priority for health and policy research. This is becoming increasingly important as life expectancies rise. Age is the biggest risk factor for AD, and lifestyle and routine health-check recommendations are in place to improve case ascertainment.^1^

The genetic epidemiology of late-onset Alzheimer’s disease (LOAD) has advanced over the last decade,^2^ with more than 20 independent loci associated with the disease in addition to *APOE.*^3^ Presently, the largest meta-analytic genome-wide association study (GWAS) for LOAD employed a two-stage study design. First, 17,008 cases were compared to 37,154 controls. 11,632 SNPs with P<1x10^−3^ from this meta-analysis were included in the second stage that compared 8,572 cases to 11,312 controls. A meta-analysis of the SNPs included in stages 1 and 2 was also performed.^4^

One difficulty and high net cost in traditional studies of AD is case ascertainment^5^ — either directly for prevalent cases or indirectly through prospective cohort studies for incident cases — given that dementia diagnosis is currently exclusively based on cognitive and functional assessment with no testing for underlying biological determinants of the clinical syndrome being required. A recent GWAS study on the UK Biobank cohort used information from family history (parent or first-degree relative with AD or dementia) as a proxy-phenotype for the participants.^6^ When meta-analysed with the GWAS summary data highlighted above,^4^ four new loci were identified.

The UK Biobank proxy-phenotype AD question, which is used here, does not incorporate biomarker data that are required for a clinical diagnosis. However, it is easy to administer at scale and we show that it has a near unit genetic correlation with the AD results from the LOAD meta-analysis^4^, where many of the samples also lacked a confirmed diagnosis by biomarker levels and autopsy.

In the present study, we related proxy-phenotype information on dementia (i.e., reporting a parent with Alzheimer’s dementia or dementia) to genetic data from 385,869 individuals from the UK Biobank cohort to identify new AD-associated loci. GWA studies were conducted separately for maternal and paternal AD due to a near two-fold difference in disease prevalence – 9.6% and 5.5%, respectively. The summary statistics from these models were meta-analysed with those from the largest publicly-available case-control study.^4^ Sensitivity analyses showed that an overlap of controls in the maternal and paternal GWAS did not bias the results. Genetic correlation analysis showed the self-reported measure of parental AD to be an accurate proxy for clinical diagnosis, validating the global meta-analysis. In addition, we tested for causal evidence of our SNP – AD associations being mediated through gene expression and DNA methylation in the prefrontal cortex.

## Subjects and Methods

### UK Biobank Cohort

UK Biobank data^7^ (http://www.ukbiobank.ac.uk) were collected on over 500,000 individuals aged between 37 and 73 years from across Great Britain (England, Wales, and Scotland) at the study baseline (2006-2010), including health, cognitive, and genetic data.

The Research Ethics Committee (REC) granted ethical approval for the study – reference 11/NW/0382 – and the current analysis was conducted under data application 10279.

### Genotyping

Genotyping details for the UK Biobank cohort have been reported previously.^8,9^ Briefly, two custom genotyping arrays were utilised with 49,950 participants typed using the UK BiLEVE Axiom Array and 438,427 participants typed using the UK Biobank Axiom Array.^8,9^ The released, genotyped data contained 805,426 markers on 488,377 individuals. Imputed genotypes were supplied with the UK Biobank data with the HRC used as the imputation reference panel.^8^

Downstream quality control steps conducted for the current analysis included removing (1) those with non-British ancestry based on both self-report and a principal components analysis, (2) outliers based on heterozygosity and missingness, (3) individuals with sex chromosome configurations that were neither XX nor XY, (4) individuals whose reported sex did not match inferred sex from their genetic data, and (5) individuals with more than 10 putative third degree relatives from the kinship table. This left a sample of 408,095 individuals. To remove the possibility of double contributions from sibs, whose parents will have the same AD status, we first considered a list of all participants with a relative (N=131,790). A genetic relationship matrix was built for these individuals using GCTA-GRM^10^ and a relationship threshold of 0.4 was applied to exclude one person from each sib-pair while retaining e.g., half-sibs and cousins and more distantly related individuals. After removing the excluded sib, the sample size was 385,869. Quality control thresholds applied to the GWAS included: minor allele frequency > 0.01, imputation quality score > 0.1, and restriction to HRC-imputed SNPs, leaving a total of 7,795,606 SNPs for the GWAS.

### Phenotypes

Family history of Alzheimer’s disease was ascertained via self-report. Participants were asked “Has/did your father ever suffer from Alzheimer’s disease/dementia?” and “Has/did your mother ever suffer from Alzheimer’s disease/dementia?” Self-report data from the initial assessment visit (2006-2010), the first repeat assessment visit (2012-2013), and the imaging visit (2014+) were aggregated with exclusions made for participants whose parents were: aged under 60 years; dead before reaching age 60 years; without age information. After merging with the genetic data, this left 32,222 cases of maternal AD with 302,756 controls, and 16,613 cases of paternal AD with 285,083 controls. Given the expected difference in disease prevalence due to sex differences in longevity – AD prevalence was double in mothers compared to fathers – GWA studies were performed separately for maternal and paternal AD.

### Genome-Wide Association Study

The GWA studies were conducted using BGENIE.^8^ The outcome variable was the residuals from a linear regression model of maternal or paternal AD on age of parent at death or at time of the offspring’s self-report, assessment centre, genotype batch, array, and 40 genetic principal components. The predictor variable was the autosomal SNP and an additive model was considered.

The GWAS linear regression coefficients were converted to odds ratios using observed sample prevalences of 0.096 and 0.055 for maternal and paternal AD, respectively,^11^ before the log-odds were multiplied by two, in this way the effect size are reported on the same scale as a traditional case-control design.^6^ Standard errors for the log-odds were then calculated based on the adjusted OR and the P-value from the initial GWAS (**Supplementary Note 1**). The ORs and standard errors were then carried forward to a weighted meta-analysis in METAL^12^ with the Stage 2 summary output from the IGAP study^4^ and the Stage 1 output for the SNPs that did not contribute to Stage 2. Linkage Disequilibrium Score (LDSC) regression was used to estimate the genetic correlation between the maternal and paternal AD GWAS results and to test for residual confounding in the meta-analysis by examining the LDSC intercept.^13,14^

The number of independent loci from the meta-analysis was determined by using the default settings in FUMA.^15^ Independent lead SNPs had P<5x10^−8^ and were independent at r^2^<0.6. Within this pool of independent SNPs, lead SNPs were defined as those in LD at r^2^<0.1. Loci were defined by combining lead SNPs within a 250kb window and all SNPs in LD of at least r^2^=0.6 with one of the independent SNPs. The 1000 genomes phase 3 data^16^ were used to map LD. This analysis was then re-run using an index SNP threshold of P<1x10^−5^ to identify suggestive loci. A gene-based analysis was carried out on all SNP output using the MAGMA software^17^ and assuming a constant sample size for all genes. A Bonferroni-adjusted P-value of 0.05/18,251 = 2.7x10^−6^ was used to identify significant genes.

### Summary-data based Mendelian Randomization

To test for pleiotropic associations between AD and gene expression/DNA methylation in the brain, summary-data based Mendelian Randomization (SMR) was performed.^18^ GWAS summary output from the meta-analysis of UK Biobank and IGAP (sample size specified as 385,869 + 74,046 = 459,915) were included along with expression QTL summary output from the Common Mind Consortium, which contains data on >600 dorsolateral prefrontal cortex samples, and DNA methylation QTL summary on 258 prefrontal cortex samples (age > 13).^19^ The reference genotypes were based on the Health and Retirement Study, imputed to the 1000 Genomes phase 1 reference panel. SNP exclusions included: imputation score <0.3, Hardy-Weinberg P-value <1x10^−6^, and a minor allele frequency <0.01. Related individuals, based on a genomic-relationship matrix cut-off of 0.05, were removed. Two sets of eQTL summary data were considered (1) after adjustment for diagnosis, institution, sex, age of death, post-mortem interval, RNA integrity number (RIN), RIN^2^, and clustered library batch (2) with additional adjustments for 20 surrogate variables. Five ancestry vectors were included as covariates in the eQTL analyses. Further details are available at: href="https://www.synapse.org/#!Synapse:syn4622659. Default parameters for the SMR analysis were used and cis eQTLs/methQTLs were considered for analysis. Bonferroni-corrected P-value thresholds were applied (P<0.05/2,011=2.5x10^−5^ for eQTL dataset 1, P<0.05/4,380=1.1x10^−5^ for eQTL dataset 2, and P<0.05/54624=9.2x10^−7^ for methQTL dataset).

## Results

### UK Biobank GWAS

There were 32,222 cases of maternal AD (302,756 controls, prevalence of 9.6%) and 16,613 cases of paternal AD (285,083 controls, prevalence of 5.5%) in UK Biobank. Linear regression GWA studies of maternal and paternal AD identified five genome-wide-significant loci, located in the *CR1*, *BIN1*, *CLU*, *PICALM*, and *APOE* gene regions (**Supplementary Figures 1 and 2**). All are established AD loci.^4^ The genetic correlation between maternal and paternal AD was not significantly different from unity (r_g_ = 0.61, SE 0.42), although the SE is large. Both traits had a high genetic correlation with the case-control summary output from the International Genomics of Alzheimer’s Disease Consortium (IGAP): r_g_ with maternal and paternal AD was 1.07 (SE 0.28) and 0.79 (0.35), respectively, both not significantly different from unity but with large SEs.

**Figure 1.**
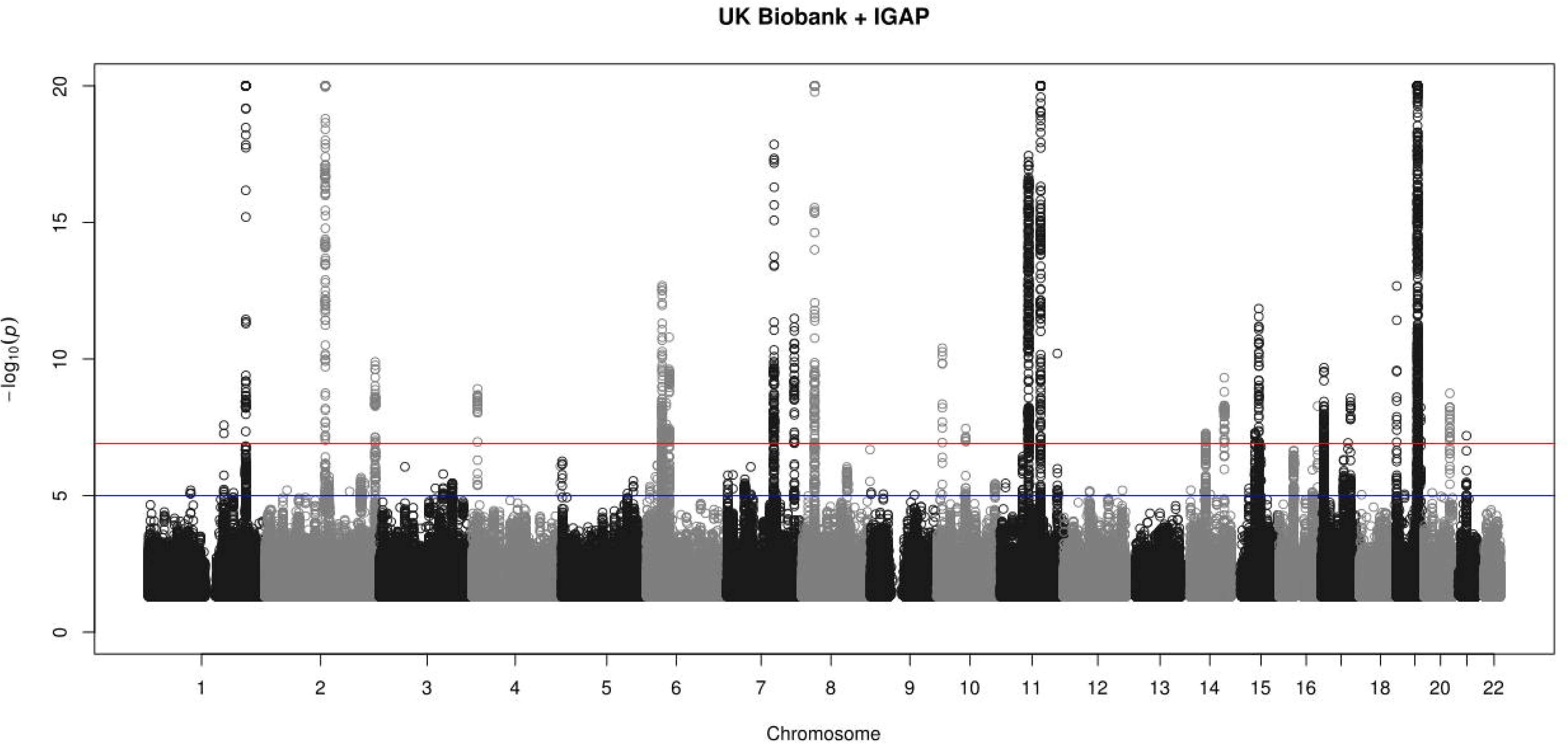
Manhattan Plot for the meta-analysis of maternal and paternal Alzheimer’s disease in UK Biobank and the results from Stage 1 and Stage 2 of IGAP [**Lambert et al. 2013**]. The red line indicates P=5x10^−8^ and the blue line indicates P=1x10^−5^. P-values truncated at 1x10^−20^

Prior to meta-analysing the UK Biobank parental summary statistics with the IGAP output, we investigated the influence of overlapping proxy-controls in UK Biobank. The p-values from a single GWAS of parental AD status (0, 1, or 2 parents with AD) correlated 0.99 with those from a meta-analysis of separate maternal AD and paternal AD; the regression of –log_10_ P-values on each other gave an intercept of 0 and a slope of 1. A meta-analysis of the summary statistics from the maternal and paternal results is therefore equivalent to the analysis of parental AD status. The linear regression effect sizes from the GWAS were converted to odds ratios prior to the meta-analysis.^11^

### Meta-Analysis

The meta-analysis of the maternal and paternal AD history in UK Biobank with the IGAP data identified 77 lead SNPs and 243 independent significant SNPs with P<5x10^−8^ from 24 genomic risk loci. The majority (n=49) of the lead SNPs were located in the gene-dense *APOE*/*TOMM40* locus on chromosome 19 (Figure 1 and **Supplementary Table 1;** GWAS summary statistics are available for the 7,795,605 meta-analysed SNPs in **Supplementary Table 2 [available online upon publication]**). The LDSC regression intercept term from the meta-analysis summary output was 1.027 (SE 0.01), indicating a polygenic signal independent from residual confounding.

**Table 1.**
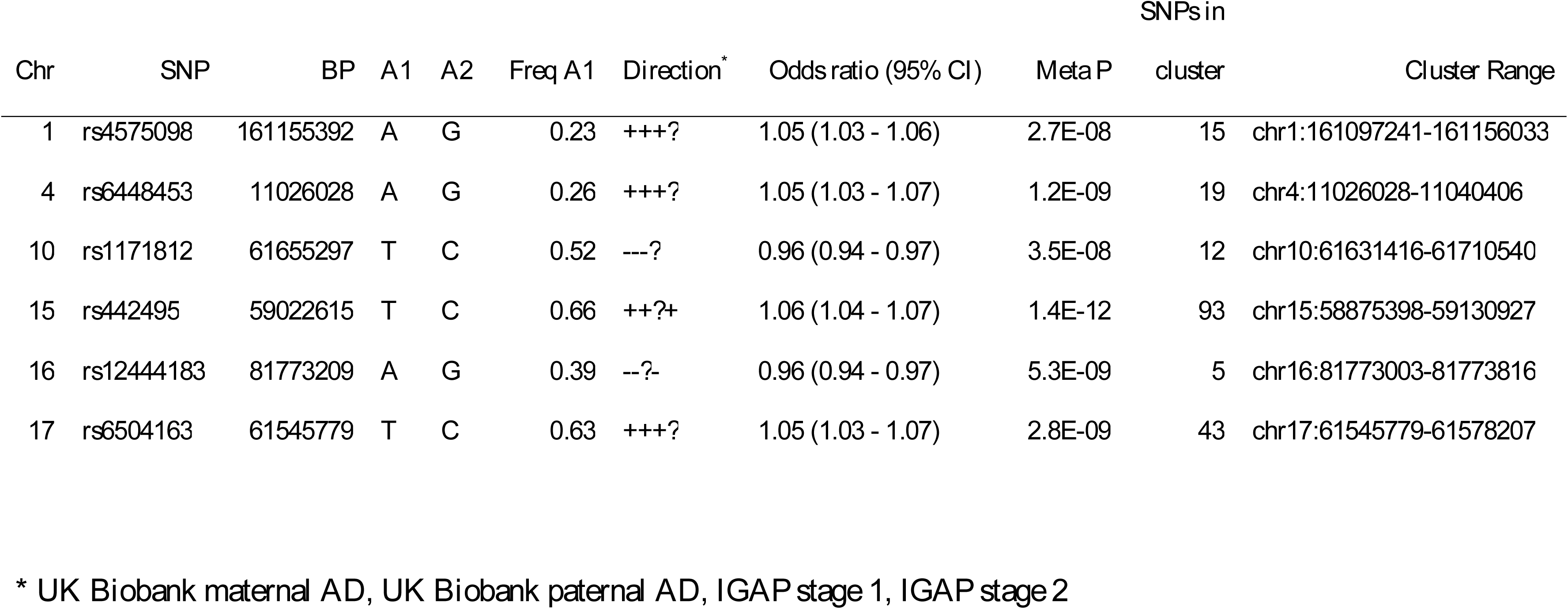
Novel SNPs (P<5x10^−8^) from the meta-analysis of UK Biobank parental history of Alzheimer’s disease with results from IGAP (**Lambert et al., 2013**).

### Novel genome-wide significant loci

Of the 24 significant risk loci, six were novel (Table 1), three of which spanned genes and gene regions with strong biological links to AD and neurodegeneration: rs4575098 (*ADAMTS4*, chr1); rs442495 (*ADAM10*, chr15); and rs6504163 (*ACE*, chr17). The other three loci had lead SNPs located in: a gene desert on chr4 (rs6448453) that is ~400kb from the *CLNK* gene (**Supplementary Figure 3)**; in *CCDC6* on chr10 (rs1171812); and in apoorly annotated region on chr16 (rs12444183), proximal to the *PLCG2* gene (**Supplementary Figure 4**). When phosphorylated, CLNK interacts with PLCG2 and is needed for PLCG2-mediated signalling in BLNK-deficient DT40 cells (http://www.uniprot.org/uniprot/Q7Z7G1#ptm_processing). PLCG2 is a transmembrane signalling enzyme important for correct functioning of the immune system.^20^ A rare variant in *PLCG2* has been found to be protective against AD.^21^

### Replication of IGAP loci

Fifteen of the 21 previously-reported SNPs^4^ associated with AD were genome-wide significant (P<5x10^−8^) in the current meta-analysis, with four other SNPs (rs2718058, rs10838725, rs17125944, and rs10498633) having P<1x10^−5^ (**Supplementary Table 3**). The *MEF2C* variant, rs190982, had a meta-analysis p-value of 5.4x10^−3^ and rs8093731 (a *DSG2* variant), which was genome-wide significant in Stage 1 but not stage 2 of IGAP, had a meta-analysis p-value of 0.18. There was complete sign-concordance between UK Biobank and IGAP for all 21 SNPs (**Supplementary Table 3**). The odds ratios between the maternal and paternal analysis for the top 21 IGAP SNPs were correlated r=0.91. Both also correlated highly with the effect sizes reported in the IGAP analysis (r = 0.85 and 0.80, respectively).

### Suggestive loci

There were 170 lead SNPs from 79 loci in the analysis considering variants with P<1x10^−5^ (Supplementary Table 4). These loci harboured SNPs that have been associated at P<5x10^−8^ with traits (Supplementary Table 5) such as: warfarin dose, triglyceride levels, BMI, Parkinson’s disease, and blood pressure (suggestive locus #64, chr16: 30,820,866-31,171,174);^22–26^ lupus and HDL cholesterol (suggestive locus #29, chr7: 50,258,234-50,318,938);^27,28^ cholesterol, heart disease, and brain white matter hyperintensity burden(suggestive locus #9, chr2: 203,639,395-204,196,618);^29–31^ asthma and allergy (suggestive locus #2, chr1: 90,302,027-90,306,216);^32^ and blood pressure (suggestive locus #68, chr17: 47,301,268-47,476,235).33).^33^ A further suggestive locus (#69, chr17: 56,398,006-56,450,524) contained a SNP within the benzodiazepine receptor (*BZRAP1*) gene that was genome-wide associated with AD in a trans-ethnic study.^34^

### Gene-based analysis

102 genes were significant at a Bonferroni threshold of P<2.7x10^−6^ (**Supplementary Table 6**). Gene Ontology analysis showed significant enrichment for the regulation of amyloid-beta clearance, negative regulation of amyloid-beta formation, very-low-density lipoprotein particle clearance, phospholipid efflux, plasma lipoprotein particle assembly, and negative regulation of endocytosis (**Supplementary Table 7**).

### Summary-data-based Mendelian Randomization (SMR)

Pleiotropic associations between AD and gene expression in the brain were tested using SMR.^18^ GWAS summary data for AD were taken from the UK Biobank and IGAP meta-analysis. eQTL summary data came from the Common Mind Consortium (n>600 dorsolateral prefrontal cortex samples: dataset 1 adjusted for age at death, sex, and institution; dataset 2 made additional adjustments for 20 surrogate variables). MethQTL data came from 258 dorsolateral prefrontal cortex samples (participants aged 13 years and older – adjustments were made for the first 5 genetic MDS components and first 11 methylation PCs).^19^ We found evidence of brain expression and DNA methylation associated with AD in the *PVR* gene (part of the *APOE*/*TOMM40* cluster on chromosome 19) in both eQTL models and also in the methQTL model (**Supplementary Tables 8-10**). However, the HEIDI p-values were <0.05 for all three analyses, indicating that the associations were unlikely to be driven by asingle causal variant affecting both expression/methylation and AD. Furthermore, different top QTL SNPs in *PVR* were identified in each of the three analyses: rs11540084, rs2301275, and rs10410915. The eQTL SNPs were in high LD in European samples^35,36^ (R^2^ = 0.99) but neither was in high LD with the methQTL (R^2^ = 0.46 and 0.45, respectively). All three SNPs are in very low LD with the *APOE* allele defining SNPs, rs7412 (max R^2^ = 0.002) and rs429358 (max R^2^ = 0.003).

## Discussion

Using recently-established proxy-phenotype methods for case ascertainment, we identified six new genome-wide significant loci for Alzheimer’s disease, three of which contain genes that have strong biological links to the disease and three others not previously linked to the disorder.

*ACE* determines levels of angiotensin II, which has trophic actions within the brain and contributes to the regulation of cerebral blood flow.^37^ Previous meta-analyses of candidate gene studies identified variants within *ACE* to be associated with AD, though not at genome-wide significance.^38,39^*ACE* variants have also been linked to atrophy of the hippocampus and amygdala,^40^ and CSF-ACE protein levels correlate with CSF tau and phosphorylated tau.^41,42^

Members of the ADAM family were identified in two of the novel loci. *ADAM10* is involved in the cleavage of amyloid beta precursor protein,^43^ which is involved in the deposition of amyloid beta, a major neurological hallmark of AD. ADAM10 has been proposed as potential therapeutic agent in AD therapy.^43,44^ Rare variants in *ADAM10* have also been linked to LOAD.^45^
*ADAMTS4* has been proposed as a regulator of synaptic plasticity during the development and ageing of the central nervous system.^46,47^

The suggestive loci (P<1x10^−5^) harboured variants in genes associated with cardiometabolic health, immunological response, and neuropathology (white matter hyperintensities and Parkinson’s disease), many of which are phenotypically linked to AD. Other suggestive loci included a *VKORC1* variant, rs9923231, whose T allele was associated with an increased risk of AD (P=2.3x10^−7^, independent SNP of locus #64), and is strongly associated with the need for a reduced dose of warfarin anticoagulation.^24,48^ The rs2526378 variant in *BZRAP1* gene (P=1.2x10^−7^, lead SNP of suggestive locus #69) was part of a cluster with rs2632516, which was a genome-wide significant SNP in a trans-ethnic study of AD.^34^
*BZRAP1* is involved in benzodiazepine receptor binding. Use of benzodiazepines has been associated with risk of dementia,^49^ particularly for users of long half-life medication^50^ thus providing a biological basis for what many considered to be an observation based on reverse causality.

An integrative analysis of eQTL and methQTL with the GWAS summary data identified one previously identified AD gene, *PVR*, as having its gene expression and methylation levels associated with AD. The most parsimonious explanation of these results is the existence of multiple causal variants, some affecting AD and others affecting expression or methylation. *PVR* is a poliovirus receptor in the *APOE*/*TOMM40* cluster on chromosome 19 that has been hypothesised to influence the risk of AD through susceptibility to viral infections.^51^ A previous SMR analysis of AD and LDL-cholesterol identified evidence of 16 pleiotropic SNPs, 12 of which were located in the *APOE* region.^52^

The main strength of the study is the proxy phenotype approach, which resulted in over 48,000 proxy-cases, which has roughly equivalent power to a study of 12,000 observed cases.^6^ However, the question used to determine parental AD status may have resulted in some responders being unable to discriminate Alzheimer’s disease and dementia from other dementia sub-types, which have different presentations and genetic architectures.^53,54^ This method of proxy-case ascertainment may have influenced the loci uncovered. Parental dementia status is partly dependent on longevity, with age being the biggest risk factor for AD. We partially controlled for this by excluding participants whose parents were younger than or died prior to reaching the age of 60 years when AD incidence is extremely low. The misclassification of case status via incorrect informant reporting will have reduced the power to detect true effects. This, along with a possible winner’s curse effect for the IGAP study, might explain the reduction in the meta-analytic odds ratios compared to those previously reported.^4^

The therapeutics field for disease modification in AD is now benefitting from more accurate, though as yet incomplete, understanding of the cascade of disease processes and phenotypic expression in preclinical and prodromal Alzheimer’s dementia populations.^55^ A prerequisite for precision medicine is the ability to identify sub-populations of a clinical condition who share a common, relevant and targetable disease mechanism.^56^ Stratification of samples for clinical trials currently being undertaken in AD rely almost exclusively on identification of intra-cerebral amyloid and therein testing of anti-amyloid therapies. Our work highlights the possibilities of basing clinical stratification on other genetic markers, in addition to *APOE*, where associated disease processes are known to be relevant, and for which pharmacological interventions are already available.

### Conclusion

We identified three new AD-associated loci that have known and putative biological processes associated with Alzheimer’s disease. Suggestive (P<1x10^−5^) associations included loci linked to common diseases and health measures that are comorbid with, or commonly used to predict risk of AD. These findings help to elucidate the biological mechanisms underlying AD and, given that some (*VKORC1*, *ACE*, *BZRAP1*) are existing drug targets for other diseases and disorders, warrant further exploration for potential precision medicine and clinical trial applications.

Supplementary information is available at MP’s website.

## Acknowledgements

Riccardo Marioni is supported by Alzheimer’s Research UK major project grant number ARUK-PG2017B-10. The research was conducted, using the UK Biobank Resource, in The University of Edinburgh Centre for Cognitive Ageing and Cognitive Epidemiology (CCACE), part of the cross-council Lifelong Health and Wellbeing Initiative (MR/K026992/1); funding from the Biotechnology and Biological Sciences Research Council (BBSRC) and Medical Research Council (MRC) is gratefully acknowledged. CCACE supports Ian Deary, with some additional support from Dementias Platform UK (MR/L015382/1). This report represents independent research part-funded by the National Institute for Health Research (NIHR) Biomedical Research Centre at South London and Maudsley NHS Foundation Trust and King’s College London. The University of Queensland authors acknowledge funding from the Australian National Health and Medical Research Council (grants 1113400, 1078901, 1078037) and the Charles & Sylvia Viertel Fellowship (JY). David Hill is supported by a grant from Age UK (Disconnected Mind Project). Alison Goate is supported by the National Institutes of Health (AG049508, AG052411) and the JPB Foundation. Alison Goate is on the scientific advisory board for Denali Therapeutics and has served as a consultant for AbbVie and Cognition Therapeutics.

The funders had no role in study design, data collection and analysis, decision to publish, or preparation of the manuscript.

## Supplemental Data Description

There are 4 supplementary figures and 10 supplementary tables

**Figure S1.** Manhattan Plot for the genome-wide association analysis of maternal Alzheimer’s disease in UK Biobank. The red line indicates P=5x10^−8^ and the blue line indicates P=1x10^−5^. P-values truncated at 1x10^−20^

**Figure S2.** Manhattan Plot for the genome-wide association analysis of paternal Alzheimer’s disease in UK Biobank. The red line indicates P=5x10^−8^ and the blue line indicates P=1x10^−5^. P-values truncated at 1x10^−20^

**Figure S3.** Plot of genome-wide significant locus #5

**Figure S4.** Plot of genome-wide significant locus #18

## Supplementary Tables (Excel file)

**Table S1.** Independent genome-wide significant AD loci from the meta-analysis of UK Biobank and IGAP summary statistics

**Table S2.** Summary output from the GWAS meta-analysis of maternal and paternal AD in UK Biobank with IGAP

**Table S3.** Lookup of genome-wide significant loci identified in Lambert et al.

**Table S4.** Independent genome-wide significant (P<5x10-8) and suggestive AD loci (P<1x10-5) from the meta-analysis of UK Biobank and IGAP summary statistics

**Table S5.** GWAS catalog output for SNPs at loci with a lead SNP at P<1x10-5

**Table S6.** MAGMA gene-based associations for the UK Biobank and IGAP meta-analysis summary results

**Table S7.** Gene Ontology enrichment analysis for the genome-wide significant genes

**Table S8.** SMR analysis of AD GWAS summary output and brain expression QTL summary output (adjusted for diagnosis, institution, sex, age of death, post-mortem interval, RNA integrity number (RIN), RIN^2, and clustered library batch)

**Table S9.** SMR analysis of AD GWAS summary output and brain methylation QTL summaryoutput (adjustment for diagnosis, institution, sex, age of death, post-mortem interval, RNA integrity number (RIN), RIN^2, clustered library batch, and 20 surrogate variables)

**Table S10.** SMR analysis of AD GWAS summary output and brain methylation QTL summary output (adjusted for 5 genetic MDS components and 11 methylation PCs)

## References

Livingston G, Sommerlad A, Orgeta V, Costafreda SG, Huntley J, Ames Det al. Dementia prevention, intervention, and care. Lancet Lond Engl 2017. doi:10.1016/S0140-6736(17)31363-6.

Huang Y, Mucke L. Alzheimer Mechanisms and Therapeutic Strategies. Cell 2012; 148: 1204–1222.

Van Cauwenberghe C, Van Broeckhoven C, Sleegers K. The genetic landscape of Alzheimer disease: clinical implications and perspectives. Genet Med Off J Am Coll Med Genet 2016; 18: 421–430.

Lambert JC, Ibrahim-Verbaas CA, Harold D, Naj AC, Sims R, Bellenguez Cet al. Meta-analysis of 74,046 individuals identifies 11 new susceptibility loci for Alzheimer’s disease. Nat Genet 2013; 45: 1452–1458.

Sibbett RA, Russ TC, Deary IJ, Starr JM. Dementia ascertainment using existing data in UK longitudinal and cohort studies: a systematic review of methodology. BMC Psychiatry 2017; 17: 239.

Liu JZ, Erlich Y, Pickrell JK. Case-control association mapping by proxy using family history of disease. Nat Genet 2017; 49: 325–331.

Sudlow C, Gallacher J, Allen N, Beral V, Burton P, Danesh Jet al. UK biobank: an open access resource for identifying the causes of a wide range of complex diseases of middle and old age. PLoS Med 2015; 12: e1001779.

Bycroft C, Freeman C, Petkova D, Band G, Elliott LT, Sharp Ket al. Genome-wide genetic data on ~500,000 UK Biobank participants. BioRxiv doi:https://doi.org/10.1101/166298.

Wain LV, Shrine N, Miller S, Jackson VE, Ntalla I, Soler Artigas Met al. Novel insights into the genetics of smoking behaviour, lung function, and chronic obstructive pulmonary disease (UK BiLEVE): a genetic association study in UK Biobank. Lancet Respir Med 2015; 3: 769–781.

Yang J, Benyamin B, McEvoy BP, Gordon S, Henders AK, Nyholt DRet al. Common SNPs explain a large proportion of the heritability for human height. Nat Genet 2010; 42: 565–569.

Lloyd-Jones LR, Robinson MR, Yang J, Visscher PM. Transformation of summary statistics from linear mixed model association on all-or-none traits to odds ratio. Genetics.

Willer CJ, Li Y, Abecasis GR. METAL: fast and efficient meta-analysis of genomewide association scans. Bioinforma Oxf Engl 2010; 26: 2190–2191.

Bulik-Sullivan B, Finucane HK, Anttila V, Gusev A, Day FR, Loh P-Ret al. An atlas of genetic correlations across human diseases and traits. Nat Genet 2015; 47: 1236–1241.

Bulik-Sullivan BK, Loh P-R, Finucane HK, Ripke S, Yang J, Schizophrenia Working Group of the Psychiatric Genomics Consortium et al. LD Score regression distinguishes confounding from polygenicity in genome-wide association studies. Nat Genet 2015; 47: 291–295.

Watanabe K, Taskesen E, Bochoven A, Posthuma D. Functional mapping and annotation of genetic associations with FUMA. Nat Commun 2017; 8: 1826.

1000 Genomes Project Consortium, Abecasis GR, Altshuler D, Auton A, Brooks LD, Durbin RM et al. A map of human genome variation from population-scale sequencing. Nature 2010; 467: 1061–1073.

de Leeuw CA, Mooij JM, Heskes T, Posthuma D. MAGMA: generalized gene-set analysis of GWAS data. PLoS Comput Biol 2015; 11: e1004219.

Zhu Z, Zhang F, Hu H, Bakshi A, Robinson MR, Powell JEet al. Integration of summary data from GWAS and eQTL studies predicts complex trait gene targets. Nat Genet 2016; 48: 481–487.

Jaffe AE, Gao Y, Deep-Soboslay A, Tao R, Hyde TM, Weinberger DRet al. Mapping DNA methylation across development, genotype and schizophrenia in the human frontal cortex. Nat Neurosci 2016; 19: 40–47.

Lehman HK. Autoimmunity and Immune Dysregulation in Primary Immune Deficiency Disorders. Curr Allergy Asthma Rep 2015; 15: 53.

Sims R, van der Lee SJ, Naj AC, Bellenguez C, Badarinarayan N, Jakobsdottir Jet al. Rare coding variants in PLCG2, ABI3, and TREM2 implicate microglial-mediated innate immunity in Alzheimer’s disease. Nat Genet 2017; 49: 1373–1384.

Warren HR, Evangelou E, Cabrera CP, Gao H, Ren M, Mifsud Bet al. Genome-wide association analysis identifies novel blood pressure loci and offers biological insights into cardiovascular risk. Nat Genet 2017; 49: 403–415.

Locke AE, Kahali B, Berndt SI, Justice AE, Pers TH, Day FRet al. Genetic studies of body mass index yield new insights for obesity biology. Nature 2015; 518: 197–206.

Takeuchi F, McGinnis R, Bourgeois S, Barnes C, Eriksson N, Soranzo Net al. A genome-wide association study confirms VKORC1, CYP2C9, and CYP4F2 as principal genetic determinants of warfarin dose. PLoS Genet 2009; 5: e1000433.

Teslovich TM, Musunuru K, Smith AV, Edmondson AC, Stylianou IM, Koseki Met al. Biological, clinical and population relevance of 95 loci for blood lipids. Nature 2010; 466: 707–713.

Nalls MA, Pankratz N, Lill CM, Do CB, Hernandez DG, Saad Met al. Large-scale meta-analysis of genome-wide association data identifies six new risk loci for Parkinson’s disease. Nat Genet 2014; 46: 989–993.

Willer CJ, Schmidt EM, Sengupta S, Peloso GM, Gustafsson S, Kanoni Set al. Discovery and refinement of loci associated with lipid levels. Nat Genet 2013; 45: 1274–1283.

Bentham J, Morris DL, Graham DSC, Pinder CL, Tombleson P, Behrens TWet al. Genetic association analyses implicate aberrant regulation of innate and adaptive immunity genes in the pathogenesis of systemic lupus erythematosus. Nat Genet 2015; 47: 1457–1464.

Dichgans M, Malik R, König IR, Rosand J, Clarke R, Gretarsdottir Set al. Shared genetic susceptibility to ischemic stroke and coronary artery disease: a genome-wide analysis of common variants. Stroke 2014; 45: 24–36.

Surakka I, Horikoshi M, Mägi R, Sarin A-P, Mahajan A, Lagou Vet al. The impact of low-frequency and rare variants on lipid levels. Nat Genet 2015; 47: 589–597.

Verhaaren BFJ, Debette S, Bis JC, Smith JA, Ikram MK, Adams HHet al. Multiethnic genome-wide association study of cerebral white matter hyperintensities on MRI. Circ Cardiovasc Genet 2015; 8: 398–409.

Pickrell JK, Berisa T, Liu JZ, Ségurel L, Tung JY, Hinds DA. Detection and interpretation of shared genetic influences on 42 human traits. Nat Genet 2016; 48: 709–717.

Newton-Cheh C, Johnson T, Gateva V, Tobin MD, Bochud M, Coin Let al. Genome-wide association study identifies eight loci associated with blood pressure. Nat Genet 2009; 41: 666–676.

Jun GR, Chung J, Mez J, Barber R, Beecham GW, Bennett DAet al. Transethnic genome-wide scan identifies novel Alzheimer’s disease loci. Alzheimers Dement J Alzheimers Assoc 2017; 13: 727–738.

Machiela MJ, Chanock SJ. LDlink: a web-based application for exploring population-specific haplotype structure and linking correlated alleles of possible functional variants. Bioinforma Oxf Engl 2015; 31: 3555–3557.

Machiela MJ, Chanock SJ. LDassoc: an online tool for interactively exploring genome-wide association study results and prioritizing variants for functional investigation. Bioinforma Oxf Engl 2017. doi:10.1093/bioinformatics/btx561.

Starr JM, Whalley LJ. ACE inhibitors: central actions. Raven Press: New York, 1994.

Elkins JS, Douglas VC, Johnston SC. Alzheimer disease risk and genetic variation in ACE: a meta-analysis. Neurology 2004; 62: 363–368.

Lehmann DJ, Cortina-Borja M, Warden DR, Smith AD, Sleegers K, Prince JAet al. Large meta-analysis establishes the ACE insertion-deletion polymorphism as a marker of Alzheimer’s disease. Am J Epidemiol 2005; 162: 305–317.

Sleegers K, den Heijer T, van Dijk EJ, Hofman A, Bertoli-Avella AM, Koudstaal PJet al. ACE gene is associated with Alzheimer’s disease and atrophy of hippocampus and amygdala. Neurobiol Aging 2005; 26: 1153–1159.

Jochemsen HM, Teunissen CE, Ashby EL, van der Flier WM, Jones RE, Geerlings MIet al. The association of angiotensin-converting enzyme with biomarkers for Alzheimer’s disease. Alzheimers Res Ther 2014; 6: 27.

Kauwe JSK, Bailey MH, Ridge PG, Perry R, Wadsworth ME, Hoyt KLet al. Genome-wide association study of CSF levels of 59 alzheimer’s disease candidate proteins: significant associations with proteins involved in amyloid processing and inflammation. PLoS Genet 2014; 10: e1004758.

Suh J, Choi SH, Romano DM, Gannon MA, Lesinski AN, Kim DYet al. ADAM10 missense mutations potentiate β-amyloid accumulation by impairing prodomain chaperone function. Neuron 2013; 80: 385–401.

Yuan X-Z, Sun S, Tan C-C, Yu J-T, Tan L. The Role of ADAM10 in Alzheimer’s Disease. J Alzheimers Dis JAD 2017; 58: 303–322.

Kim M, Suh J, Romano D, Truong MH, Mullin K, Hooli Bet al. Potential late-onset Alzheimer’s disease-associated mutations in the ADAM10 gene attenuate {alpha}-secretase activity. Hum Mol Genet 2009; 18: 3987–3996.

Krstic D, Rodriguez M, Knuesel I. Regulated proteolytic processing of Reelin through interplay of tissue plasminogen activator (tPA), ADAMTS-4, ADAMTS-5, and their modulators. PloS One 2012; 7: e47793.

Lemarchant S, Pomeshchik Y, Kidin I, Kärkkäinen V, Valonen P, Lehtonen Set al. ADAMTS-4 promotes neurodegeneration in a mouse model of amyotrophic lateral sclerosis. Mol Neurodegener 2016; 11: 10.

Wadelius M, Chen LY, Downes K, Ghori J, Hunt S, Eriksson Net al. Common VKORC1 and GGCX polymorphisms associated with warfarin dose. Pharmacogenomics J 2005; 5: 262–270.

Starr JM, Whalley LJ. Drug-induced dementia. Incidence, management and prevention. Drug Saf 1994; 11: 310–317.

Shash D, Kurth T, Bertrand M, Dufouil C, Barberger-Gateau P, Berr Cet al. Benzodiazepine, psychotropic medication, and dementia: A population-based cohort study. Alzheimers Dement J Alzheimers Assoc 2016; 12: 604–613.

Porcellini E, Carbone I, Ianni M, Licastro F. Alzheimer’s disease gene signature says: beware of brain viral infections. Immun Ageing 2010; 7: 16.

Zhu Z, Zheng Z, Zhang F, Wu Y, Trzaskowski M, Maier R et al. Causal associations between risk factors and common diseases inferred from GWAS summary data. bioRxiv 2017;: 168674.

Ferrari R, Wang Y, Vandrovcova J, Guelfi S, Witeolar A, Karch CMet al. Genetic architecture of sporadic frontotemporal dementia and overlap with Alzheimer’s and Parkinson’s diseases. J Neurol Neurosurg Psychiatry 2017; 88: 152–164.

Ikram MA, Bersano A, Manso-Calderón R, Jia J-P, Schmidt H, Middleton Let al. Genetics of vascular dementia - review from the ICVD working group. BMC Med 2017; 15: 48.

Ritchie CW, Molinuevo JL, Truyen L, Satlin A, Van der Geyten S, Lovestone Set al. Development of interventions for the secondary prevention of Alzheimer’s dementia: the European Prevention of Alzheimer’s Dementia (EPAD) project. Lancet Psychiatry 2016; 3: 179–186.

Hampel H, O’Bryant SE, Castrillo JI, Ritchie C, Rojkova K, Broich Ket al. PRECISION MEDICINE - The Golden Gate for Detection, Treatment and Prevention of Alzheimer’s Disease. J Prev Alzheimers Dis 2016; 3: 243–259.

